# BK Channel Function Bidirectionally Influences Behavioral Deficits Following Withdrawal from Chronic Ethanol in *Caenorhabditis elegans*

**DOI:** 10.1101/062752

**Authors:** Luisa L. Scott, Scott J. Davis, Rachel C. Yen, Greg J. Ordemann, Deepthi Bannai, Jonathan T. Pierce-Shimomura

**Author notes:** To whom correspondence should be addressed: Jonathan T. Pierce-Shimomura, University of Texas at Austin, Neuroscience Department, 2506 Speedway NMS 5.234, Mailcode C7350, Austin, TX 78712.

## Abstract

The severity of withdrawal from chronic ethanol is a driving force for relapse in alcohol dependence. Thus, uncovering molecular changes that can be targeted to lessen withdrawal symptoms is key to breaking the cycle of dependence. Using the model nematode *Caenorhabditis elegans*, we tested whether one highly conserved ethanol target, the BK potassium channel, may play a major role in alcohol withdrawal. Consistent with a previous report, we found that *C. elegans* displays behavioral impairment during withdrawal from chronic ethanol that can be reduced by low-dose ethanol. We discovered that the degree of impairment is exacerbated in worms lacking the BK channel, SLO-1, and is alleviated by selective rescue of the BK channel in the nervous system. Conversely, behavioral impairment during withdrawal was dramatically lower in worms with BK channel function enhanced via gain-of-function mutation or overexpression. Consistent with these results, we found that chronic ethanol exposure decreased BK channel expression in a subset of neurons. In addition, we found that a distinct, conserved large-conductance potassium channel, SLO-2, showed the inverse functional relationship, influencing withdrawal behavior via a SLO-1 channel-dependent mechanism. Our findings demonstrate that withdrawal symptoms in *C. elegans* are mechanistically explained in part by a functional imbalance in the nervous system associated with a reduction in SLO-1 channel expression. Therefore, selective modulation of Slo family ion channel activity may represent a novel therapeutic approach to explore for normalizing behaviors during alcohol withdrawal.

**ARTICLE SUMMARY:** People addicted to alcohol maintain maladaptive drinking patterns in part to avoid the terrible symptoms of withdrawal. It is unclear whether any single molecule may be genetically modified to alleviate withdrawal symptoms. Here, we discover that for the nematode *C. elegans*, upregulating function of the conserved BK potassium channel SLO-1 prevents alcohol withdrawal symptoms. Conversely, downregulating SLO-1 channel function makes withdrawal worse. Moreover, we identify an inverse relation between SLO-1 and a second type of BK channel named SLO-2 in the severity of withdrawal. The BK channel thus represents an attractive molecular target to consider for alleviating alcohol withdrawal symptoms.

**Statement on data and reagent availability:** Strains are available upon request or through the *Caenorhabditis* Genetics Center.

## INTRODUCTION

Neural adaptation during persistent exposure to ethanol underlies many of the symptoms of withdrawal from chronic alcohol consumption (Koob *et al.* 1998; Koob 2013). These symptoms include life-threatening conditions such as seizures and rapid heart rate as well as psychological conditions such as anxiety and confusion (Finn and Crabbe 1997). The severity of symptoms, particularly the degree of negative affect, following withdrawal from chronic ethanol is a driving force for relapse (Winward *et al.* 2014). Uncovering targets that modulate the neural state in withdrawal to more closely match the naïve state is important for developing pharmacological agents that will ameliorate withdrawal symptoms and thus reduce relapse (Becker and Mulholland 2014). While many cellular proteins are implicated as mediators of the effects of ethanol *in vivo*, a single target has not yet been found to dominate control over withdrawal behavior in any species.

The large-conductance, calcium-and voltage-activated BK potassium channel (Slo1) is a well-conserved target of ethanol across species as diverse as worm, fly, mouse and man (Mulholland *et al.* 2009; Treistman and Martin 2009; Bettinger and Davies 2014). Clinically relevant concentrations (10-100 mM) of ethanol are sufficient to alter Slo1 gating in *in vitro* preparations (Chu and Treistman 1997; Jakab *et al.* 1997; Dopico *et al.* 1998, 2003; Walters *et al.* 2000; Brodie *et al.* 2007). Impairing Slo1 function *in vivo* influences ethanol behaviors, such as acute intoxication and tolerance (Davies *et al.* 2003; Cowmeadow *et al.* 2005, 2006; Martin *et al.* 2008; Kreifeldt *et al.* 2013). Prolonged ethanol exposure further changes the expression of Slo1 channels, suppressing overall levels and replacing channels with ethanol-insensitive isoforms (Pietrzykowski *et al.* 2008; Velazquez-Marrero *et al.* 2011; Li *et al.* 2013; N’Gouemo and Morad, 2014). This responsiveness to ethanol has generated interest in the Slo1 channel as a potential target for influencing the severity of withdrawal from ethanol (Ghezzi *et al.* 2012; N’Gouemo and Morad 2014). Recent experiments with mouse knockouts of proteins that modulate Slo1 function (that is, non-essential auxiliary subunits) suggest this channel alters escalation of drinking in a withdrawal paradigm (Kreifeldt *et al.* 2013). However, testing a direct role of these channels in withdrawal has been precluded by the behavioral and physiological deficits exhibited by Slo1 knock out mice (e.g. Thorneloe *et al.* 2005; Meredith *et al.*2006; Pyott *et al.* 2006; Typlt *et al.* 2013; Lai *et al.* 2014).

To surmount these pleitrophic deficits of the Slo1 KO mouse and directly probe whether Slo1 function contributes to behavioral deficits during alcohol withdrawal, we used the tiny nematode, *Caenorhabditis elegans*. Previously, unbiased forward genetic screens demonstrated the importance of neuronal expression of the worm ortholog of the Slo1 channel, called SLO-1, for acute ethanol intoxication in naïve worms (Davies *et al.* 2003). Ethanol activated the native worm SLO-1 channel in neurons at the same concentration (20-100 mM) as previously shown for human Slo1 channels (Davies *et al.* 2003). Loss-of-function mutations in *slo-1* rendered worms resistant to intoxication, while gain-of-function mutations in *slo-1* caused worms to appear intoxicated in the absence of alcohol (Davies *et al.* 2003). Here we show that SLO-1 function has the opposite relation for alcohol withdrawal. Chronic alcohol appears to lower SLO-1 function which contributes to behavioral impairments during alcohol withdrawal. The extent of withdrawal-induced impairments is worse in the absence of SLO-1, while increasing activity or overexpression of SLO-1 improves withdrawal behaviors. Consistent with previous findings in mammalian cells *in vitro* (Pietrzykowski *et al.* 2008; Ponomarev *et al.* 2012; N’Gouemo and Morad 2014), SLO-1 expression declined in some neurons during chronic ethanol exposure *in vivo*. Another member of the large conductance potassium-channel family, SLO-2, which is gated by intracellular calcium and/or chloride (Yuan *et al.* 2000; Zhang *et al.* 2013), showed a relationship to alcohol withdrawal that was inverse to that of the SLO-1 channel; behavioral impairment during withdrawal was stemmed by null mutations in the *slo-2* gene. Together, our results suggest that Slo channels are part of the neural adaptation to chronic ethanol exposure in *C. elegans*, and that increasing SLO-1 channel activity, either directly or indirectly, rebalances neural circuits underlying these behaviors.

## Materials and Methods

### Animals

*C. elegans* strains were grown at 20 °C and fed OP50 strain bacteria seeded on <u>N</u>ematode <u>G</u>rowth <u>M</u>edia (NGM) agar plates as described previously (Brenner 1974). Worms cultured on plates contaminated with fungi or other bacteria were excluded. The reference wild-type (WT) strain was N2 Bristol. The background for the *slo-1(null)* rescue strains was NM1968, harboring the previously characterized null allele, *js379* (Wang *et al.* 2001). The background *slo-1(null);slo-2(null)* double mutant strain was JPS432, obtained by crossing NM1968 with LY100 and confirmed via sequencing. This latter strain harbored the previously characterized *slo-2* null allele, *nf100* (Santi *et al.* 2003). Strains NM1630 and LY101 were used as *slo-1(null)* and *slo-2(null)* reference strains, respectively. JPS1 carried the previously characterized *slo-1* gain-of-function allele, *ky399* (Davies *et al.* 2003).

### Transgenesis

Multi-site gateway technology (Invitrogen, Carlsbad, CA) was used to construct plasmids for the *slo-1* rescue and overexpression strains. 1894 kb of the native *slo-1* promoter (*pslo*-1) was used to drive *slo-1a(cDNA)::mCherry-unc-54* UTR expression. *punc-119* was used as a pan-neuronal promoter (Maduro and Pilgrim 1995). All plasmids were injected at a concentration of 20 ng/μL for rescue for in a *slo-1(null)* background and 5-10 ng/μL for overexpression in a WT background (Mello *et al.* 1991). The co-injection reporter PCFJ90 *pmyo-2:mCherry* (1.25 ng/μl) was used to ensure proper transformation of the arrays. Two independent isolates were obtained for most strains to help control for variation in extrachromosomal arrays. The following strains were generated for this study: JPS344 *slo-1(null) vxEx344* [*pslo-1::slo-1a::mCherry::unc-54UTR pmyo-2::mCherry*], JPS345 *slo-1(null) vxEx344* [*pslo-1::slo-1a::mCherry::unc-54UTR* + *pmyo-2::mCherry*], JPS529 *slo-1(null) vxEx529* [*punc-119::slo-1a::mCherry::unc-54UTR* + *pmyo-2::mCherry*], JPS523 *slo-1(null);slo-2(null) vxEx523* [*pslo-1::slo-1a::mCherry::unc-54UTR* + *pmyo-2::mCherry*], JPS524 *slo-1(null);slo-2(null) vxEx524* [*pslo-1::slo-1a::mCherry::unc-54UTR* + *pmyo-2::mCherry*], JPS521 *vxEx521* [*pslo-1::slo-1a::mCherry::unc-54UTR* + *pmyo-2::mCherry*] (injected at 5 ng/μL), JPS522 *vxEx522* [*pslo-1::slo-1a::mCherry::unc-54UTR* + *pmyo-2::mCherry*] (injected at 10 ng/μL). To image mCherry-tagged SLO-1 protein expression, we used strains JPS572 *slo-1(null);vsIs48* [*punc-17::GFP*] *vxEx345* [*pslo-1::slo-1a::mCherry::unc-54UTR* + *pmyo-2::mCherry*], and JPS576 *slo-1(null);slo-2(null) vxEx576* [*pslo-1::slo-1a::mCherry::unc-54UTR* + *pmyo-2::mCherry* + *prab-3::GFP*]. To determine whether the *slo-1* promoter was sensitive to chronic ethanol treatment, we used strain JPS584 *vxEx584* [*pslo-1(rescue)::GFP::unc-54UTR* + *ptph-1::mCherry*].

### Ethanol treatment

Methods for assaying ethanol withdrawal were modified from Mitchell *et al.* (2010). Well-populated (>200 worms), 6-cm diameter plates were bleached to obtain eggs, which were allowed to grow age-matched to the mid- to-late-stage L4-larval stage. L4 worms derived from the same plate were then divided between an ethanol-infused (+ethanol) and standard control (−ethanol) seeded plate. Standard plates were 6-cm diameter Petri dishes filled with 12-mL NGM-agar and seeded with OP50 bacteria. Ethanol plates (400 mM) were prepared by adding 280 μL of 200-proof ethanol (Sigma Aldrich) beneath the agar of the standard seeded plates and allowing the ethanol to soak into the agar. The plates were sealed with Parafilm and worms were exposed for 20-24 hours. The ethanol-treated worms were withdrawn on standard seeded plates for one hour. Worms kept on the standard seeded plate overnight served as the naive controls.

### RNAi treatment

RNA interference (RNAi) was performed as previously described (Timmons and Fire 1998; Fraser *et al.* 2000). In brief, RNAi plates were prepared by adding IPTG (1 mM) and carbenicillin (25 μg/mL) to standard unseeded plates. These plates were seeded with bacterial strain HT115 transformed with vector L4440 containing genomic fragments to the *slo-2* gene (*F08B12.3*, Source Bioscience). Worms were moved to the RNAi plates at mid-to late L4-larval stage.

### Diacetyl race assay

Methods were modified from Bargmann *et al.* (1993) and Mitchell *et al.* (2010). Race plates were prepared by drawing a start and a goal line on the bottom of standard unseeded, 6-cm diameter Petri dishes filled with 12-mL NGM-agar. The race plates were prepared within 20 minutes of each race by applying a 10-μL mixture of attractant (1:1000 dilution of diacetyl) and paralytic (100-mM sodium azide) at the goal. Worms were cleaned of bacteria by transferring them to one or more unseeded plates until they left no residual tracks of bacteria, a process that took less than 10 minutes. Approximately 25 worms were placed on the start side of the race plate by transferring with a platinum pick and the total number of worms, and the number of worms that reached the goal, were counted every 15 minutes for one hour to calculate the percent of worms at the goal. Counts were performed with the observer blind to genotype and experimental treatment.

### Locomotion assay

Worms were cleaned of bacteria as described above and approximately 15 were moved into a 5/8-inch diameter copper ring sealed on a standard unseeded plate (see above). Movement was recorded for 2 minutes at 2 frames/second with a FLEA digital camera (Point Grey, Richmond, BC, Canada). The distance that the worms crawled during one minute was measured using a semi-automated procedure in ImagePro Plus (Media Cybernetics, Rockville, MD) to objectively calculate overall speed of individual worms.

### Gas chromatography

Internal ethanol measurements were estimated using previous methods (Alaimo *et al.* 2012). Only a fraction of the external ethanol enters worms when treated on NGM agar plates; but see Mitchell *et al.*, 2007 for an alternate view of how ethanol enters worms incubated in liquid Dent’s medium. For WT worms, we measured the internal ethanol concentration at 0, 20 min, 3 hours and 24 hours of ethanol treatment as well as after 1 hour of withdrawal. For other strains, the internal ethanol concentration was measured at 24 hours and one hour after withdrawal. Worms exposed to ethanol as described above were rinsed with ice-cold NGM buffer into a 1.5-mL Eppendorf tube and briefly spun (< 10 sec) at low speed to separate the worms from the bacteria. The liquid was removed, replaced with ice-cold NGM buffer and the sample was spun again. All of the liquid was carefully removed to leave only the worm pellet. This pellet was then doubled in volume with ice-cold NGM buffer. The sample went through five rapid freeze-thaw cycles using liquid nitrogen plus 30 seconds of vortexing and was finally spun down at high speed for 2 minutes. Two microliters of the sample was added to a gas chromatography vial. The amount of ethanol was measured using headspace solid-phase microextraction gas chromatography (HS-SPME-GC). Automation of the HS-SPME-GC measurement was obtained using an autosampler (Combi Pal-CTC Analytics, Basel, Switzerland). Ethanol analysis was carried out using a gas chromatograph equipped with a flame ionization detector.

### Confocal microscopy

First-day adult worms were mounted on 2% agarose pads, immobilized with 30mM sodium azide and imaged with a Zeiss laser-scanning microscope (LSM710) using Zen (black edition) acquisition software (Carl Zeiss, Germany). GFP fluorescence and phase contrast images were collected using a 488-nm laser and mCherry fluorescence was collected using a 561-nm laser. Once set, the laser power and electronic gain were held constant for the red and green channels to perform ratiometric analysis. Using a 63X water immersion objective and a 0.9-micron pinhole, neurons were imaged in three dimensions taking slices every 0.8 microns through the z-axis. Ratiometric analysis was completed in ImageJ (Schneider *et al.* 2012). Z-stacks through the neurons were summed, and the mean pixel intensity was measured for the red and green channel in the area of interest. Background intensity was measured using the same size region of interest next to the worm. This background measurement was then subtracted from the neuronal measurement.

### Statistical analysis

Sigmaplot 12.5 (Systat Software, San Jose, CA) was used for all statistical analyses to determine significance (p ≤ 0.05, two tailed) between two or more groups. Groups were compared using t- or ANOVA tests where appropriate. If needed, post hoc multiple comparisons were performed using the Holm-Sidak method. All measures were obtained with the observer blind to genotype and experimental treatment.

## RESULTS

### Behavioral deficits during withdrawal recovered by low-dose ethanol

To test how *C. elegans* behaves during withdrawal from chronic ethanol exposure, we modified a treatment paradigm based on Mitchell *et al.* (2010). In brief, wild-type (WT), age-matched, L4-stage larvae were treated with ethanol for 24 hours and then withdrawn for 1 hour on seeded control plates (red timeline in Figure 1a, see Methods for details). A control naïve group of worms was set up in parallel (black timeline in Figure 1a). We used gas chromatography to estimate the worms’ internal ethanol concentration at 0, 20 min, 3 hours and 24 hours of ethanol treatment, as well as after 1 hour of withdrawal. Internal ethanol concentration rose gradually to ~50 mM over 3 hours, consistent with non-instantaneous uptake of the ethanol from the agar substrate (Figure 1b).*C. elegans* only absorbs a fraction of the high external concentration of ethanol (400 mM) when assayed on standard plates (Alaimo *et al.* 2012). Internal ethanol concentration remained at ~50 mM after 24-hour exposure and returned to baseline values after withdrawal (Figure 1b).

**Figure 1.**
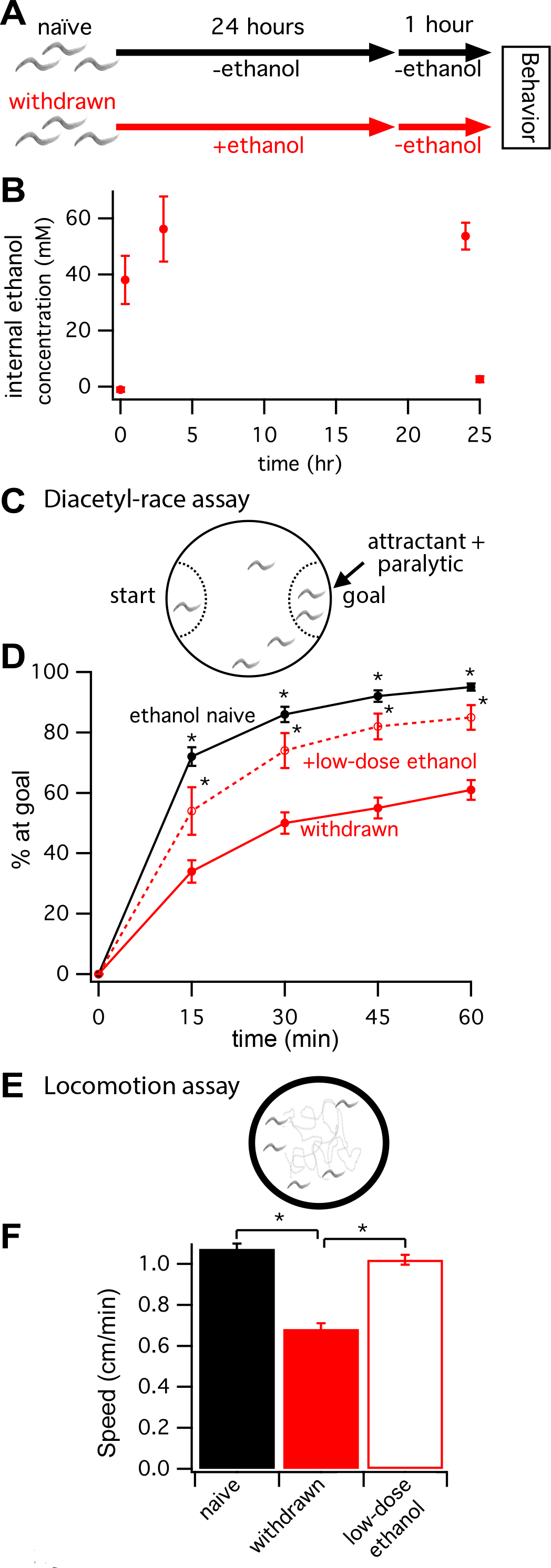
Two behavioral deficits during alcohol withdrawal recovered by low-dose ethanol. Worms withdrawn from chronic ethanol exposure display behavioral deficits. (A) Schematic showing the exposure paradigm used for the two treatment groups, naive (black) and withdrawn (red), starting with age-matched L4-stage larvae. Worms assayed for behaviors are young adults 25 hours later. (B) Gas chromatography determined internal ethanol concentration after 0, 20 min, 3 hours and 24 hours of ethanol treatment, and after 1 hour of withdrawal. (C) Schematic of the diacetyl-race assay. Diacetyl was used as a volatile attractant and sodium azide was used as a paralytic trapping worms that reached the goal. (D) The mean percent of worms that reached the attractant +/− SEM plotted every 15 minutes for 1 hour. At all timepoints, withdrawn worms (solid red line) performed less well than naive worms (solid black line). Withdrawn worms treated with a low dose of ethanol during the race (dashed red line) performed significantly better than withdrawn worms (* p < 0.05). (E) Schematic of locomotion assay. Worms were allowed to move freely on a blank agar surface within a copper ring. (F) Histogram of mean speed +/− SEM. Locomotion was also impaired during withdrawal. Withdrawn worms moved slower than naive worms (naive vs. withdrawn, 1.10 ± 0.026 vs. 0.68 ± 0.028 cm/min, p < 0.001). Again, this withdrawal-induced impairment was improved when worms were placed on low-dose ethanol during the assay (withdrawn vs. +low-dose ethanol, 0.68 ± 0.028 vs. 1.0 ± 0.025 cm/min, p < 0.001).

Next, we assayed the behavioral performance of worms in a chemotaxis race to the attractant diacetyl (Figure 1c). Specifically, we calculated the percent of worms that migrated across a 6-cm diameter, unseeded plate to reach a spot of diacetyl at the goal every 15 minutes for 1 hour. At a 1:1000 dilution, diacetyl is a strong volatile chemoattractant (Bargmann *et al.* 1993). Worms were paralyzed at the goal by the addition of sodium azide to the chemoattractant solution (Figure 1c). Withdrawn worms and ethanol-naïve controls from the same age-matched cohort were raced in tandem on different plates. Similar to findings by Mitchell *et al.* (2010) for performance in a 10-cm food-race assay, we found that worms withdrawn from chronic ethanol treatment showed impaired diacetyl-race performance relative to untreated, ethanol-naïve worms (Figure 1d). The performance of withdrawn worms improved on race plates with a low dose of exogenous ethanol (less than 25% of the ethanol concentration during the 24 hour exposure, Figure 1d).

In separate assays without diacetyl, we determined that basal locomotion was also impaired during withdrawal. Crawling on unseeded plates (Figure 1e) was ~40% slower for withdrawn worms than naive worms (naive vs. withdrawn, 1.10 ± 0.026 vs. 0.68 ± 0.028 cm/min, p < 0.001; Figure 1f). Again, this withdrawal-induced impairment was improved when worms were treated with low-dose ethanol (withdrawn vs. withdrawn + low-dose ethanol, 0.68 ± 0.028 vs. 1.0 ± 0.025 cm/min, p < 0.001; Figure 1f). Thus, in agreement with Mitchell *et al.* (2010), we find that *C. elegans* displays the fundamental traits of alcohol withdrawal symptoms observed in higher animals including humans, i.e. behaviors are impaired after removal from a prolonged exposure to ethanol, and these impairments can be partly rectified by re-exposure to a low dose of ethanol.

### Withdrawal impairments worsened by reduced neuronal SLO-1 channel function

To ascertain whether these behavioral impairments during withdrawal hinge on changes in SLO-1-channel activity or expression, we looked at withdrawal behavior in a number of strains with altered *slo-1* expression. Withdrawn performance was assessed as a function of naïve performance to account for any baseline effects of the genetic modifications. These effects were small unless otherwise noted (see Figure S1). Two *slo-1* null alleles, *js379* and *js118*, showed significantly stronger withdrawal-related impairment on the diacetyl-race assay than the WT strain N2 (Figure 2a; *js379* vs. N2, p < 0.05; *js118* vs. N2, p < 0.005). These same strains also showed greater withdrawal-induced slowing in locomotion than WT (Figure 2b; *js379* vs. N2, p < 0.05; *js118* vs. N2, p < 0.01). These impairments were rescued by extrachromosomal expression of *slo-1(+)* in a *slo-1(null)* background using different promoters. Rescue strains with *slo-1(+)* driven by the endogenous promoter *(pslo-1)* or a pan-neuronal promoter *(punc-119)* showed substantially improved performance on the diacetyl-race assay compared to the background *slo-1(js379)* null strain (Figure 2a; versus *slo-1(null)*, p < 0.001). Intriguingly, the *slo-1(+)* pan-neuronal rescue strain performed the diacetyl-race assay at a level equivalent to naïve WT worms (Figure 2a). Rescue strains with *slo-1(+)* driven by either promoter also displayed less impaired locomotion upon withdrawal compared to the background *slo-1(js379) null* strain (Figure 2b). The deleterious effect of deleting *slo-1* on withdrawal behaviors was not due to differences in ethanol uptake or metabolism. A strain with the canonical *slo-1(js379)* null allele showed no difference in internal ethanol concentration at 24 hours of ethanol treatment or after one hour of withdrawal than WT worms (Figure S2; *slo-1(null)* vs. N2, n.s.).

**Figure 2.**
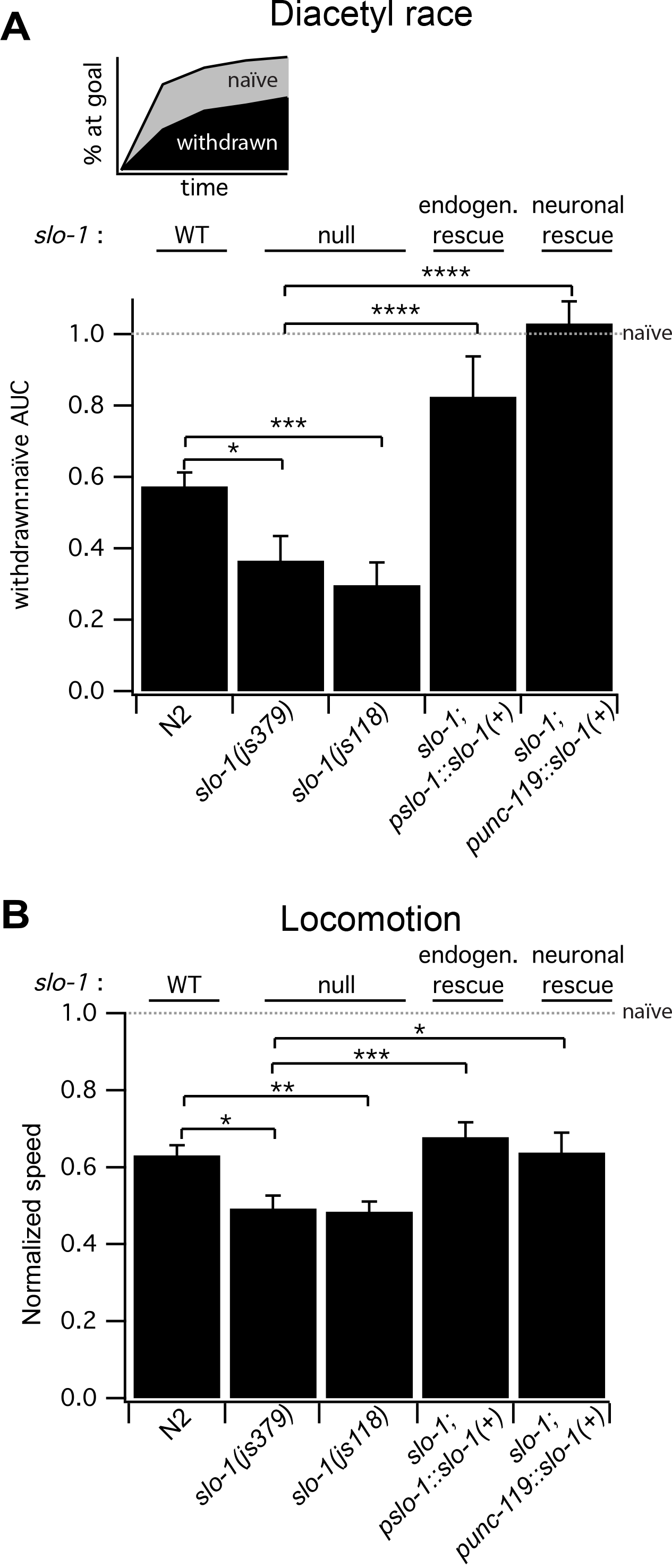
Reduced neuronal SLO-1 channel function exacerbated behavioral impairments during alcohol withdrawal. (A) Schematic above indicates how the time course of performance was quantified by the <u>a</u>rea <u>u</u>nder the <u>c</u>urve (AUC) for the percent of worms at the goal vs. time for the diacetyl race. Treatment groups: withdrawn (black area), naïve (gray + black areas). Histogram below shows the mean AUC for withdrawn worms normalized to the mean AUC for naive worms (dashed horizontal line) +/− SEM. The *slo-1* genotype for each strain is indicated above each bar for reference. Two *slo-1* strains with null alleles (*js379* and *js118*) were more impaired upon withdrawal for the diacetyl-race assay than WT strain N2. Rescue strains with *slo-1(+)* driven by the endogenous promoter (*pslo-1*) or a pan-neuronal promoter (*punc-119*) showed substantially improved performance on the diacetyl-race assay compared to the background *slo-1* null mutant. (B) Locomotion during withdrawal also worsened with reduced BK channel function. Histogram shows mean speed during withdrawal for different strains normalized to mean speed for naive worms (dashed horizontal line) +/− SEM. Two *slo-1* null strains were more impaired upon withdrawal for locomotion than WT. Rescue strains with *slo-1(+)* driven by the endogenous promoter or a pan-neuronal promoter showed substantially improved performance compared to the background null strain. For panels A and B, * p < 0.05, *** p < 0.005, and **** p < 0.001.

### Withdrawal impairments improved by enhancing SLO-1 channel expression or activity

Thus far our findings showed that reducing SLO-1 channel expression in neurons exacerbated withdrawal-induced behavioral impairments. Next, we ascertained whether increasing SLO-1 channel function could improve withdrawal behaviors. A previously characterized gain-of-function allele, *slo-1(ky399)*, showed significantly less impairment in the diacetyl-race assay upon withdrawal (Figure 3; *slo-1(ky399)* vs. N2, p < 0.001). Independent overexpression strains made by transforming with low (5 ng/mL) or moderate (10 ng/mL) *slo-1(+)* DNA in a WT background also showed less withdrawal-induced impairment in the diacetyl-race assay (Figure 3a; both strains versus N2, p < 0.001). Finally, the *slo-1* gain-of-function and *slo-1(+)* overexpression strains also showed less impairment in locomotion upon withdrawal (Figure 3b; all three strains versus N2, p < 0.05).Just as for the *slo-1* null strains, differences in ethanol uptake or metabolism did not appear to account for the protective effect of enhancing SLO-1 function on withdrawal behavior (Figure S2; *slo-1* overexpression strain vs. N2, n.s.). Together, these data suggested that increasing BK channel activity reduces withdrawal-induced behavioral deficits.

**Figure 3.**
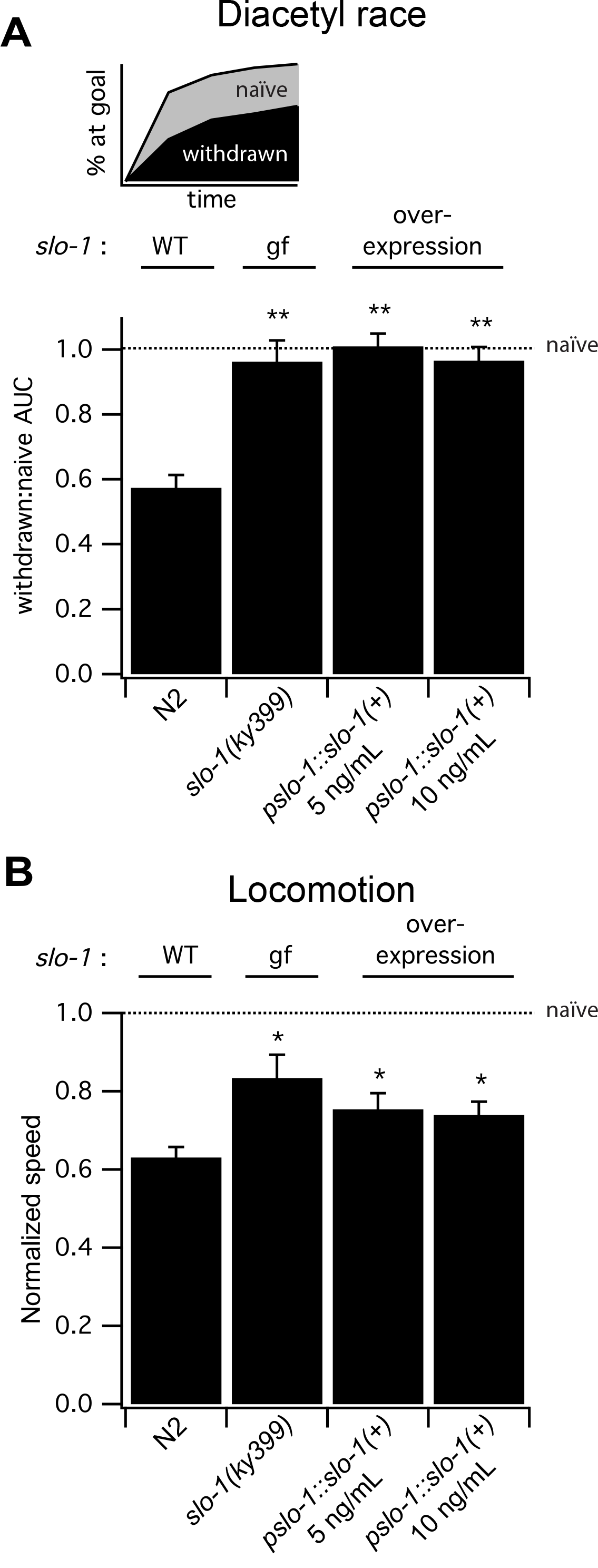
Enhanced SLO-1 channel function ameliorated behavioral impairment during alcohol withdrawal. (A) Schematic above indicates how performance was quantified by the are<u>a</u>rea <u>u</u>nder the <u>c</u>urve (AUC) for the percent of worms at the goal vs. time for the diacetyl race. Treatment groups: withdrawn (black area), naïve (gray + black areas). Histogram below shows the mean AUC for withdrawn worms normalized to the mean AUC for naïve worms (dashed horizontal line) +/− SEM. The *slo-1* genotype for each strain is indicated above each bar for reference. The *slo-1(ky399)* gain-of-function mutant and two strains with *slo-1 (+)* overexpressed in a WT background were significantly less impaired upon withdrawal for the diacetyl-race assay than WT strain N2. (B) Enhancing SLO-1 channel function also improved locomotion during withdrawal. Histogram shows mean normalized crawl speed +/− SEM. For panels A and B, * p < 0.05, *** p < 0.005, and **** p < 0.001.

Interestingly, locomotion both in naïve and withdrawn worms was dependent upon non-neuronal expression of *slo-1* (e.g., muscle) and/or seemingly precise levels of neuronal expression. Pan-neuronal expression of *slo-1(+)* with the *punc-119* promoter was sufficient to normalize diacetyl-race performance in naive worms (Figure S1a). However, naïve worms with pan-neuronal expression of *slo-1(+)* displayed slow locomotion (Figure S1b). Withdrawn locomotion for this strain was only comparatively better than the background strain (Figure 2b,Figure S1b). Locomotion appeared to be sensitive to too much SLO-1 expression or activity, even under the endogenous *pslo-1* promoter. A *slo-1* overexpression strain transformed with 10 ng/μL, but not 5 ng/μL, showed impaired locomotion under naive conditions. A strain carrying a *slo-1* gain-of-function allele performed particularly poorly under naïve conditions. These strains with poor naive performance had comparatively, but not absolutely, faster crawl speeds during withdrawal than WT (Figure 3b). Thus, although different approaches of enhancing BK channel function uniformly improved withdrawal behaviors, they had more variable effects on the behaviors, particularly crawling, of naïve worms.

### SLO-2, a distinct large-conductance potassium channel, influences withdrawal impairments via a SLO-1 channel-dependent mechanism

Concerted regulation of the activity or tone of multiple ion channels by changes in neuronal activity supports homeostatic function of the nervous system following perturbations (O’Leary *et al.* 2014). Our behavioral genetic results suggest that Slo1 channel function is down regulated with chronic alcohol exposure, and that this underlies the behavioral impairments during withdrawal. Like mammals, worms have more than one large-conductance potassium channel in the Slo family, specifically SLO-1 and SLO-2 (Yuan *et al.* 2000; Santi *et al.* 2003). The SLO-2 channel was recently shown to carry a large portion of outward rectifying current in many worm neurons (Liu *et al.* 2014). Recent evidence suggests that, like SLO-1, *C. elegans* SLO-2 is activated by intracellular Ca^2^+ and voltage (Zhang *et al.* 2013) suggesting that SLO-2 could play a similar role in neuronal function as SLO-1 in worm. Co-expression and co-regulation in sensory neurons suggest that these channels could act in concert to regulate behavior (Alqadah *et al.* 2016). Accordingly, we tested whether blocking SLO-2 function influenced withdrawal behavior. This was assayed using the diacetyl race because this paradigm better probed the role of neuronal *slo-1* expression in ethanol withdrawal (see above). In contrast with our findings for SLO-1, we found that two independent *slo-2* null alleles, *nf100* and *nf101*, showed improved performance in the diacetyl-race assay during withdrawal from chronic ethanol relative to WT (Figure 4a; *nf100* and *nf101* vs. N2, p < 0.001). In addition, we found that WT worms treated with *slo-2* RNAi from L4 larval-stage onward also displayed less behavioral impairment during withdrawal (Figure 4a; RNAi treated N2 vs. control N2, p < 0.001). This suggests that improved diacetyl racing of *slo*-2*(null)* mutants during withdrawal may be accounted for by even a relatively short-term down regulation of *slo-2* after most of the nervous system has already developed by the L4 larval stage.

**Figure 4.**
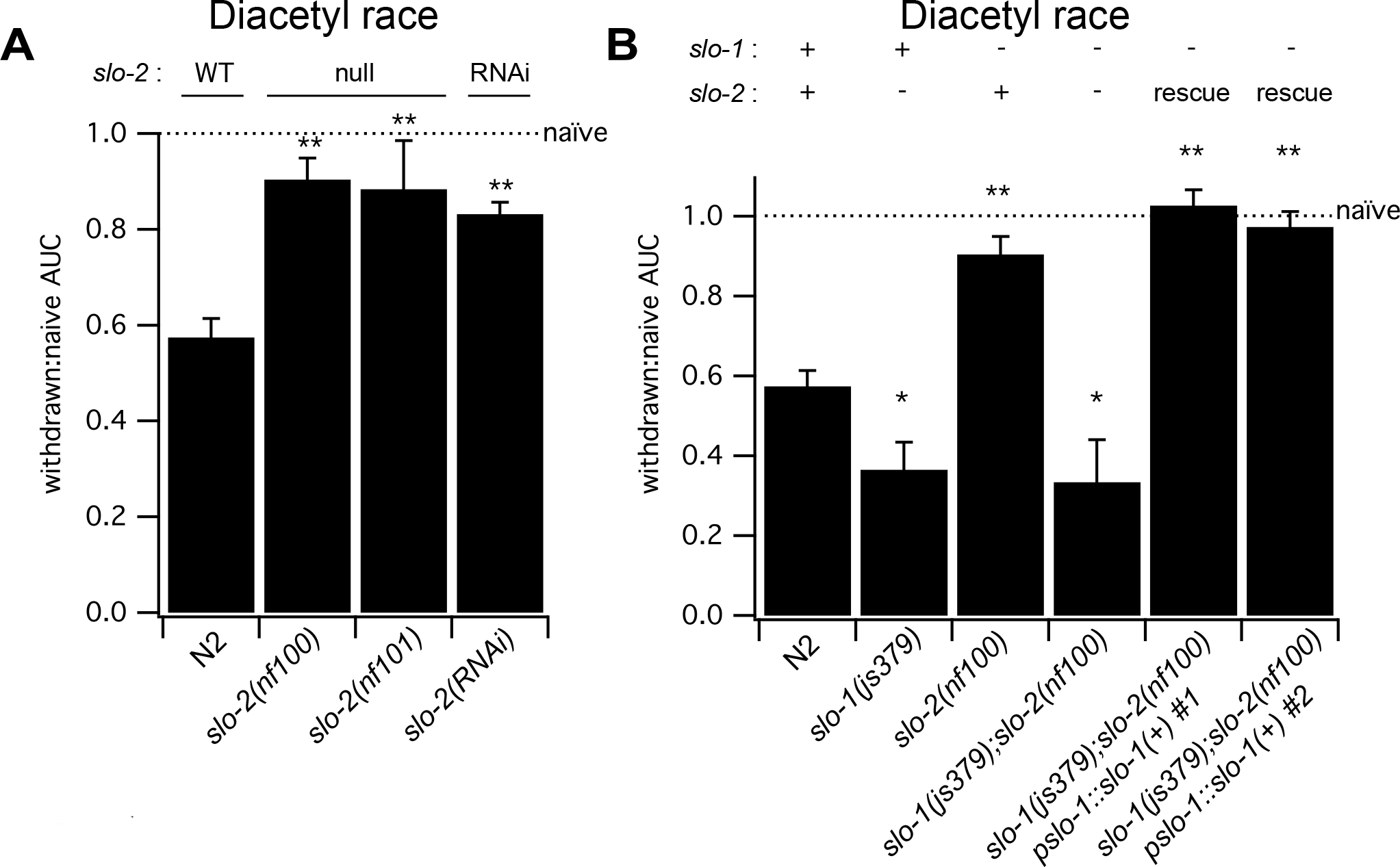
A different large-conductance potassium channel, SLO-2, influences withdrawal impairments via a SLO-1 channel-dependent mechanism. Knock-out of *slo-2* improved behavior during alcohol withdrawal.(A) Histogram shows the mean <u>a</u>rea <u>u</u>nder the <u>c</u>urve (AUC) values of different strains for diacetyl-race performance; withdrawn performance normalized to naïve performance (dashed lines) +/− SEM. Two *slo-2* null strains were significantly less impaired upon withdrawal for the diacetyl race than WT. Similarly, WT worms treated with *slo-2* RNAi during EtOH treatment showed less behavioral impairment during withdrawal. (B) Epistasis between *slo-1* and *slo-2* for alcohol withdrawal. A strain carrying null alleles for both *slo-1* and *slo-2* was more impaired in the diacetyl race during withdrawal than WT, similar to the parent *slo-1* null strain. Independent rescue strains (#1 and #2) with *slo-1(+)* introduced on the *slo-1;slo-2* double null mutant background were less impaired than the parent stain during withdrawal. For panels A and B, *p < 0.025 and **p < 0.001.

We next performed epistasis analysis to probe the genetic relation between *slo-1* and *slo-2* in alcohol withdrawal. Although the *slo-2* null allele *nf100* alone conferred resistance to withdrawal-induced behavioral deficits, when both *slo-1* and *slo-2* genes were knocked out, the resulting double mutant was more impaired in the diacetyl race during withdrawal than WT, similar to the parent *slo-1(js379)* null mutant (Figure 4b; *slo-1(js379);slo-2(nf100)* vs. N2, p < 0.025; *slo-1(js379);slo-2(nf100)* vs. *slo-1(js379)*, n.s.). The protective effects of the *slo-2* null allele against withdrawal-induced impairments returned when *slo-1(+)* was reintroduced under the endogenous promoter (Figure 4b; both rescue strains versus *slo-1(js379);slo-2(nf100)*, p < 0.001). Finally, the protective effect of the *slo-2* null allele did not appear to be due to differences in ethanol uptake or metabolism. The *slo-2* null mutant showed no difference in internal ethanol concentration at 24 hours of ethanol treatment or after one hour of withdrawal compared to WT worms (Figure S2). Together these results showed that knocking out the SLO-2 channel protects against withdrawal-induced behavioral impairments. Moreover, this protection is conferred via a *slo-1*-mediated mechanism.

### Chronic ethanol treatment suppresses SLO-1 channel expression in some neurons

Our behavioral genetic evidence for reduced SLO-1 channel function lead us to predict that chronic ethanol treatment may suppress neuronal Slo1 channel expression as observed in vertebrates (Pietrzykowski *et al.* 2008; N’Gouemo and Morad 2014). Because previous analysis found no difference in whole-worm levels of *slo-1* transcripts in naïve and ethanol-treated conditions (Kwon *et al.* 2004), we chose to investigate differences in the expression of SLO-1 protein. We used the endogenous promoter for *slo-1* (pslo-1) to express mCherry-tagged SLO-1 protein in specific neurons for naïve and chronic ethanol-treated worms.

The *slo-1(js379)* null background was used to minimize functional consequences of overexpressing the *slo-1* transgene. The amount of red fluorescence was expressed as a function of GFP-labeling in representative neurons that participate in locomotion (VC4 and VC5 motor neurons) or in odor sensation (left or right AWA sensory neuron pair) behaviors (Vidal-Gadea *et al.* 2011; Faumont *et al.* 2011; Bargmann *et al.* 1993). The red:green fluorescence ratio decreased by half in motor neurons after the 24-hour ethanol exposure (Figure 5a, p < 0.0001), but showed no significant change in sensory neurons (Figure 5b). These findings suggest that SLO-1 channel expression levels may be decreased, but not abolished, by ethanol exposure in a subset of neurons important for locomotory aspects of the diacetyl race and locomotion performance.

**Figure 5.**
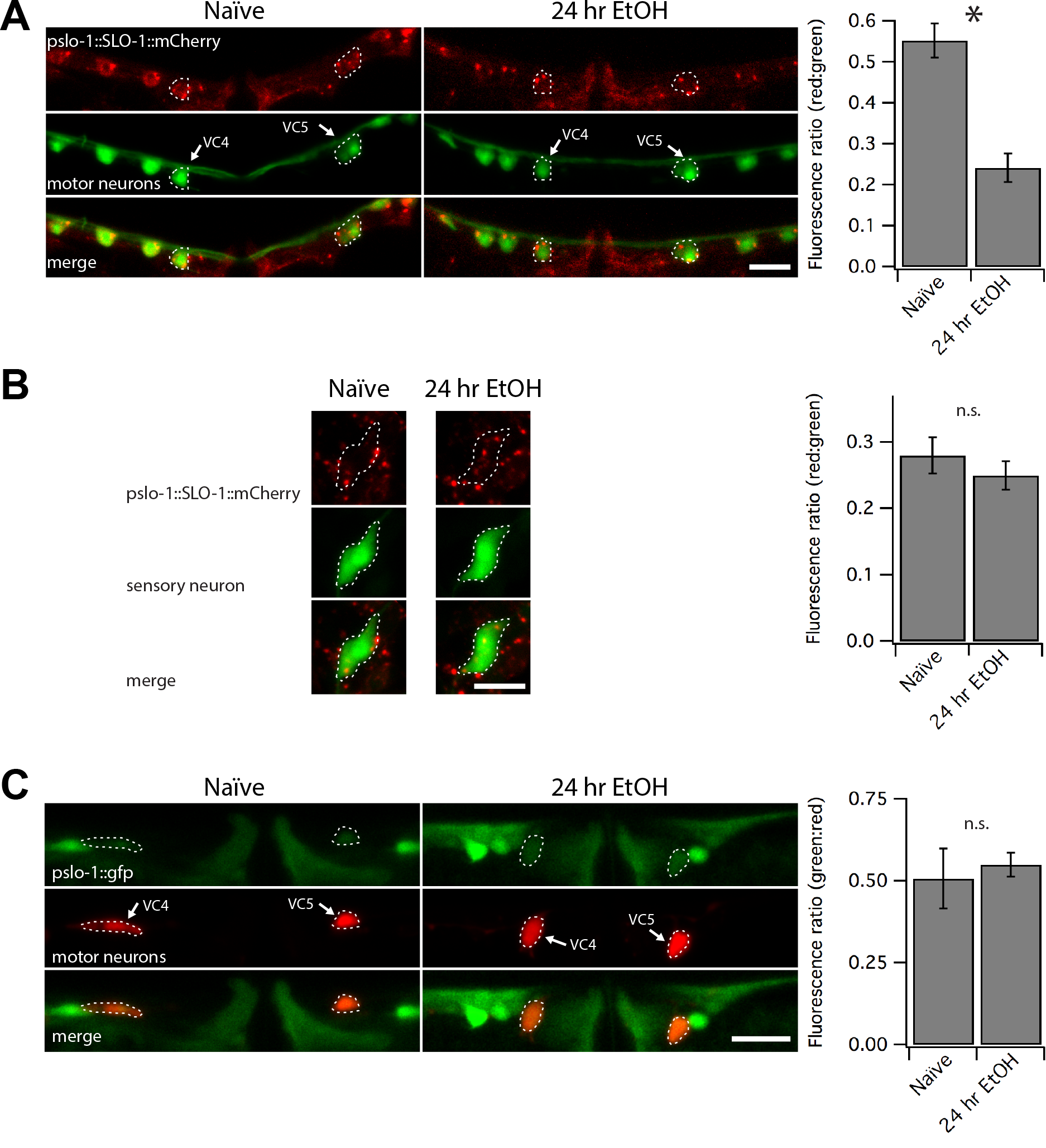
Chronic ethanol treatment suppresses neuronal SLO-1 channel expression. (A,B) Confocal microscopy stacks were summed to produce the photomicrographs showing translational *slo-1* reporter tagged with mCherry in a *slo-1(null)* background. The red:green fluorescence decreased by half in GFP-labeled VC4 and VC5 neurons after 24-hour exposure to ethanol (A, *** p < 0.0001), but not in GFP-labeled AWA olfactory neurons (B). (C) Confocal photomicrographs showing a GFP transcriptional reporter of *slo-1* in the green channel and mCherry-labeled VC4 and VC5 motor neurons. Ratiometric analysis showed no change in whole body green:red ratios in the VC4 and VC5 neurons following chronic ethanol treatment. Scale bars represent 10 μm in panels A-C.

To further probe the mechanisms by which SLO-1 channel expression is downregulated by chronic ethanol exposure, we tested whether the *slo-1* promoter has ethanol-sensitive elements by fusing the same slo-1 promoter (pslo-1) to generate a *pslo-1:gfp* transcriptional reporter in a WT background. This promoter region was sufficient to rescue or improve behavioral phenotypes above. To perform ratiometric analysis, the transcriptional reporter was coexpressed on the same extrachromosomal array with a second reporter that labels VC4 and 5 motor neurons with mCherry under a *ptph-1* promoter that was previously shown to be ethanol insensitive (Kwon *et al.* 2004). Expression of the transcriptional reporter was not altered in VC4 and VC5 motor neurons in response to 24 hours of ethanol exposure (Figure 5c). We conclude that the decrease in mCherry-tagged SLO-1 channel after chronic ethanol exposure may be due to post-translational processes.

## DISCUSSION

Here we show that worms withdrawn from chronic ethanol exposure displayed behavioral deficits suggestive of altered nervous system function. Chronic ethanol has been found to change many aspects of nervous system function and whole body physiology in animal models including gene expression in worm (e.g. Lovinger and Roberto 2013; Nagy, 2004; Osterndorff-Kahanek *et al.* 2015; Kwon *et al.* 2004). Nevertheless, we found that simply increasing SLO-1 channel tone, even selectively in neurons, was sufficient to overcome these behavioral deficits in *C. elegans.* Conversely, we found that the extent of withdrawal-induced impairments was far worse in the absence of SLO-1 channels. This bidirectional relation between SLO-1 channel function and withdrawal severity appears to be explained in part by a decrease in SLO-1 channel function over prolonged exposure to ethanol. A decrease in SLO-1 channel function may represent an adaptive response to the activating effect that ethanol has on the SLO-1 channel (Davies *et al.* 2003). Adaptive modulation of SLO-1 channel tone, either through functional changes or channel expression, may also be linked to the activity of other ion channels; we discovered that the extent of withdrawal-induced impairment was modulated oppositely by a second highly conserved member of the large conductance potassium family, SLO-2, via a *slo-1*-dependent mechanism. These results suggest that Slo1 may represent a molecular target to alleviate withdrawal symptoms in higher animals.

### Withdrawal as a neuroadaptive response to prolonged ethanol exposure

Many studies support the theory that alcohol abuse disorders including addiction are accompanied by, or even may be caused by, adaptive responses by the nervous system to chronic alcohol consumption (Koob 2013; Koob 2015).Initially, prolonged exposure to ethanol triggers homeostatic changes to nervous system function that act to maintain near normal function. Over time, particularly when ethanol is removed from the system, some of these homeostatic changes may lead to pathological dysfunction, contributing to alcohol disorders.

Our results are consistent with the idea that Slo1 expression is regulated as part of the neural adaptation to chronic ethanol exposure. Acutely, ethanol acts directly to modulate the function of the Slo1 channel (Dopico *et al.*, 2016). In *C. elegans*, ethanol increases the open probability of the SLO-1 channel both *in vivo* and *in vitro* (Davies *et al.* 2003; Davis *et al.* 2015). Long-term, homeostatic down regulation of Slo1 channel function would compensate for hyperactivation of the Slo1 channel in the presence of ethanol. Once the worm is removed from ethanol, however, the hypothetical homeostatic decrease in SLO-1 channel tone would contribute to dysfunctional behaviors. This may account for why behavioral deficits induced by withdrawal from ethanol could be overcome with either overexpression of the SLO-1 channel or gain-of-function mutation in the SLO-1 channel. The severe withdrawal symptoms of the *slo-1* mutant worms may be similarly explained if knock-out of *slo-1* overcompensates the homeostatic reduction of SLO-1 channel tone.

### Mechanisms for neuroadaption to chronic ethanol

How might Slo1 function be lowered during chronic ethanol exposure? We found that for *C. elegans*, one way chronic ethanol appears to down regulate Slo1 channel tone is to reduce expression in select neurons. This is based on ratiometric analysis of a reduction in mCherry-tagged SLO-1 channels in the soma of certain motor neurons but not sensory neurons. The SLO-1 channel is widely expressed throughout the nervous system and muscle (Wang *et al.* 2001). Adaptive neuronal changes in SLO-1 expression may only occur in some neurons, perhaps to normalize the function of circuits that depend on a critical level of SLO-1 activity for behaviors. We did not observe a corresponding decrease in the expression of a *slo-1* transcriptional reporter in the same neurons, consistent with a previous report showing no overall ethanol-induced downregulation of *slo-1* transcription in whole worms nor evidence of a consensus sequence for an ethanol responsive element in the *slo-1* promoter (Kwon *et al.* 2004). Instead, we propose that the reduction in SLO-1 expression may occur at the protein level. This could occur via post-translational modification of processes controlling SLO-1 degradation or distribution (reviewed in Kyle and Braun 2014). For example, a ubiquitination pathway is used to regulate Slo1 expression in response to seizures. Following seizure activity Slo1 is ubiquitinated and retained in the endoplasmic reticulum (Liu *et al.* 2014) where it presumably enters into the endoplasmic reticulum-associated degradation pathway. A similar mechanism may decrease Slo1 expression to normalize circuit activity in the face of chronic ethanol exposure.

Slo1 function is regulated multiple ways by ethanol in other systems. Previous work showed that 24 hours of ethanol exposure *in vitro* was sufficient to downregulate expression of Slo1 transcripts in mammalian neurons (Peitryzykowski *et al.* 2008). Whole-cell Slo1 currents are also downregulated in rat neurons during withdrawal (N’Gouemo and Morad 2014). Interestingly, alcohol-dependent humans were shown to have a downregulation of Slo1 transcripts in frontal cortex (Ponomarev *et al.* 2012). Further balancing the effect of ongoing ethanol activation of Slo1 channels, the fraction of ethanol-insensitive isoforms increases with ethanol exposure in mammals (Peitryzykowski *et al.* 2008, Gouemo and Morad 2014; Velázquez-Marrero *et al.* 2011). In addition, the Slo1 channel function is readily altered at the post-translational level by kinases and other signaling pathways influenced by ethanol (Shipston and Tian 2016; Dopico *et al.* 2014; Ron and Jurd 2005).

Given this evidence for varied modulation of Slo1 channel function by ethanol in other systems, we suspect that SLO-1 function is also down regulated with chronic ethanol exposure via multiple mechanisms in worm. For instance, it remains to be tested if ethanol exposure alters the expression profile of the ten *slo-1* isoforms in *C. elegans* (Johnson *et al.* 2011; LeBoeuf and Garcia 2012), like Slo1 splice variation is influenced by ethanol in mammalian cells (Peitryzykowski *et al.* 2008). Based on the subtly of our finding of differential SLO-1 regulation in specific neurons, future studies investigating transcriptional regulation of *slo-1* by ethanol will need to somehow 1) differentiate transcripts from the adult nervous system versus those from other tissues and the developing worms harbored in eggs within the adult, and 2) differentiate expression changes between neurons and within neuron compartments. Additionally, given the importance of splice variation in Slo1 expression, function and sensitivity to ethanol (Shipston and Tian 2016; Dopico *et al.* 2014), a complete investigation of the translational modification of the SLO-1 channel in worm in the future will need to consider all aspects of endogenous SLO-1 regulation. While transcriptional changes in splice isoforms are also likely, the reduced expression of mCherry-tagged SLO-1 encoded by *slo-1* cDNA we find here specifically points to post-translational mechanisms for ethanol down regulation of SLO-1 expression in a subset of worm neurons.

Intriguingly, we found that a second highly conserved member of the large-conductance potassium family, SLO-2, also modulates neural adaptation during chronic ethanol exposure and subsequent withdrawal. This modulation occurs via a *slo-1*-dependent mechanism. Mammalian Slo2 channels are expressed in neurons where they influence action potential propagation and shape synaptic integration (Bhattacharjee and Kaczmarek 2005). Worm SLO-1 and SLO-2 exhibit a similar expression profile as in mammal, both in neurons and muscle influencing multiple behaviors, and share a similar means of activation, along with co-expression and co-regulation (Zhang *et al.* 2013; Alqadah *et al.* 2016). These findings suggest that in worm SLO-1 and SLO-2 channels could act in concert to regulate behavior. For example, SLO-1 and SLO-2 show redundant regulation of the terminal fate of asymmetric sensory neurons in worm (Alqadah *et al.* 2016); however, SLO-2 but not SLO-1 regulates hypoxia (Zhang *et al.* 2013). Here we show another interaction between these channels with anticorrelated regulation of alcohol withdrawal. Thus, deletion of *slo-2* may be compensated by increased expression or altered composition of SLO-1, or changes in shared signaling pathways that enhance SLO-1 channel gating. We speculate that the resistance to behavioral impairment due to alcohol withdrawal in *slo-2* mutants may be explained by an adaptive process of enhanced SLO-1 function, particularly evident after exposure to ethanol.

In addition to the *slo-1* mutants described here, two other mutants defective in neuromodulatory signalling were previously found to be required for ethanol withdrawal phenotypes (Mitchell *et al.* 2010). The first mutant, *npr-1*, is missing a worm ortholog to the vertebrate neuropeptide-Y receptor, while the second mutant, *egl-3*, is missing a propeptide convertase that is required for cleavage of hundreds of neuropeptides (Mitchell *et al.* 2010). Such signaling has already been implicated in ethanol responses in mammals. It will be interesting to find out whether *slo-1* represents a major downstream target in these two neuromodulatory pathways during alcohol withdrawal.

### Slo1 plays a central role in responses to ethanol across behaviors

Previously, through two large, independent, unbiased forward-genetic screens, the *slo-1* gene encoding the SLO-1 channel was found to represent the most important single gene required for acute intoxication in *C. elegans* (Davies *et al.* 2003). Our findings here show that the SLO-1 channel also plays a central, but opposite role in neuronal plasticity during alcohol withdrawal in worm. Analogous opposite short and long term functional roles of the Slo1 channel in alcohol behaviors may be expected in higher animals.

## Acknowledgments

Support for this study was provided by a NRSA award F31AA021641 to S. J. D by NIAAA, as well as the Waggoner Center, ABMRF, NIAAA R03AA020195, and R01AA020992 to J. T. P.-S. We thank the *Caenorhabditis* Genetic Center (funded by the NIH), Drs. Hongkyun Kim and Ikue Mori for reagents, as well as Susan Rozmiarek for expert assistance.

---

Supplementary Figure 1. **Direct comparison of performance during withdrawal.** (A) Mean percent of worms at goal plotted every 15 minutes for withdrawn worms for the diacetyl race. Key for different strains on right near corresponding traces. Error bars +/− SEM for WT strain N2, but other error bars omitted for clarity. This includes separate mean data plotted for pairs of distinct overexpression, *slo-1(+)* rescue, and *slo-1* null strains. (B) Bars show mean speed during withdrawal +/− SEM for strains with various altered *slo-1.* OE means overexpression in WT background. Endogenous promoter is *pslo-1* and panneuronal promoter is *punc-119*. (C) Bars shows mean speed during withdrawal +/-SEM for strains with mutant *slo-2*.

Supplementary Figure 2. **Altering SLO-1 or SLO-2 channels did not appear to cause differences in ethanol uptake or metabolism.** (A) Mean internal ethanol concentration after 24-hour exposure to ethanol measured by gas chromotagraphy in WT strain N2, *slo-1(js379)* null mutant, *slo-1(+)* overexpression and *slo-2(nf100)* null mutant strains +/− SEM. There was no significant difference in concentration between these strains. (B) Mean internal ethanol concentration for the same strains after 1 hour of withdrawal from ethanol +/−SEM. There was no significant difference in concentration between strains.

